# Impact of Capillary and Sarcolemmal Proximity on Mitochondrial Structure and Energetic Function in Skeletal Muscle

**DOI:** 10.1101/2024.01.08.574684

**Authors:** Hailey A. Parry, T. Bradley Willingham, Kevin A. Giordano, Yuho Kim, Shureed Qazi, Jay R. Knutson, Christian A. Combs, Brian Glancy

**Affiliations:** National Lung, Blood, and Heart Institute, National Institutes of Health, Bethesda, MD, USA; Shephard Center’s Virginia C. Crawford Research Institute, Atlanta, GA, USA; Holy Cross Orthopedic Institute, Fort Lauderdale, FL, USA; University of Massachusetts, Lowell, MA,USA; National Institute of Arthritis and Musculoskeletal and Skin Diseases, National Institutes of Health, Bethesda, MD, USA

## Abstract

Mitochondria within skeletal muscle cells are considered to be located either between the muscle contractile apparatus (interfibrillar mitochondria, IFM) or in large pools beneath the cell membrane (subsarcolemmal mitochondria, SSM), with several structural and functional differences reported between IFM and SSM. However, recent 3D imaging studies suggest that proximity to capillaries embedded in sarcolemmal grooves, rather than proximity to the sarcolemma itself, may drive the accumulation of mitochondria near the cell periphery (paravascular mitochondria, PVM). To evaluate the impact of capillary versus sarcolemmal proximity, we compared the structure and function of skeletal muscle mitochondria located either in large pools lateral to embedded capillaries (PVM), adjacent to the sarcolemma but not in PVM pools (SSM), or interspersed between sarcomeres (IFM). Mitochondrial morphology and interactions were assessed by 3D electron microscopy coupled with machine learning segmentation while mitochondrial energy conversion was assessed by two-photon microscopy of mitochondrial membrane potential, content, calcium, NADH redox and flux in live, intact cells. Structurally, while PVM and SSM were similarly larger than IFM, PVM were more compact and had greater mitochondrial connectivity compared to both IFM and SSM. Functionally, PVM had similar or greater basal NADH flux compared to SSM and IFM, respectively, despite a more oxidized NADH pool and a greater membrane potential, signifying a greater activation of the electron transport chain in PVM. Together, these data indicate proximity to capillaries has a greater impact on mitochondrial energy conversion and distribution in skeletal muscle than the sarcolemma alone.

## Introduction

Mitochondria in striated muscles were first observed 180 years ago (Schwann, 1838; Henle, 1841) and later identified to be critical for energy production (ATP) and metabolic regulation (Claude, 1940; Lardy & Wellman, 1952, 1953; Keilin & King, 1958; Mitchell, 1961; Ernster & Schatz, 1981). Early imaging studies of striated muscle revealed cell-type specialization of mitochondrial localization when examining the subcellular localization of mitochondria in oxidative and glycolytic muscle cells types. Specifically, initial 2D light microscopy images discovered that mitochondria in oxidative fibers are primarily localized to subsarcolemmal regions surrounding myonuclei and directly parallel to the sarcomeric A-bands, whereas mitochondrial aligned perpendicular to the sarcomeric I-bands in glycolytic fibers (Kölliker, 1857; Cajal, 1888; Glancy & Balaban, 2021). Based on these initial observations and subsequent electron microscopy studies (Romanul, 1964, 1965; Bubenzer, 1966b; Gauthier & Padykula, 1966; Hoppeler *et al*., 1973; Bakeeva *et al*., 1978b; Kirkwood *et al*., 1986a), striated muscle mitochondria are often classified as either subsarcolemmal mitochondria (SSM) or interfibrillar mitochondria (IFM) despite forming a continuous network (Bakeeva *et al*., 1978b; Kirkwood *et al*., 1986a; Glancy *et al*., 2015). The SSM are considered those which cluster beneath the sarcolemma (cell membrane) and IFM are found interspersed between myofibrils (Bubenzer, 1966b; Hoppeler *et al*., 1973; Bakeeva *et al*., 1978b; Kirkwood *et al*., 1986a; Picard *et al*., 2012). Notably, these region-specific mitochondrial pools differ in composition, structure, and function, along with distinct responses to both exercise and disease (Krieger *et al*., 1980; Cogswell *et al*., 1993; Bizeau *et al*., 1998; Adhihetty *et al*., 2005; Koves *et al*., 2005; Ritov *et al*., 2005; Ferreira *et al*., 2010; Nielsen *et al*., 2010; Chomentowski *et al*., 2011; Kavazis *et al*., 2017).

Because organelle function is intrinsically linked to structure, it is important to consider how structural difference among SSM and IFM pools may contribute to the subcellular specialization of mitochondrial function. As first observed with 2D light and electron microscopy, IFM are long, thin projections of mitochondria due to their location between myofibrils (Romanul, 1964, 1965; Gauthier & Padykula, 1966; Hoppeler *et al*., 1973). SSM, in contrast, are considered large, globular organelles due, in part, to the lack of spatial constraints imposed by the contractile apparatus (Rothstein *et al*., 2005; Picard *et al*., 2012; Picard *et al*., 2013a; Picard *et al*., 2013b; Vincent *et al*., 2019; Willingham *et al*., 2021). However, the 2D images from which these mitochondrial localization models are built upon often lack the critical subcellular context of other nearby, but out of frame, structures which may impact mitochondrial morphology and function. Indeed, 3D light microscopy analyses of the mouse Tibialis anterior muscle *in vivo* suggested that the large pools of mitochondria located beneath the sarcolemma in oxidative fibers were always associated with a nearby capillary and that the volume of mitochondria within these regions was directly related to the depth to which capillaries were embedded in grooves in the sarcolemma (Glancy *et al*., 2014). Based on these data, we hypothesized that proximity to the capillary, rather than sarcolemma, drives the observed pooling of mitochondria near the cell periphery (Glancy & Balaban, 2021; Willingham *et al*., 2021). Therefore, we anticipate that systematic evaluation subcellular mitochondrial heterogeneity will reveal three, rather than two, regional classifications of skeletal muscle mitochondria: mitochondria in large pools lateral to the capillary (PVM), mitochondria directly under the sarcolemma but not within the PVM (SSM), and mitochondria interspersed among the myofibrils (IFM; Figure 1). However, to date, there have been no quantitative assessments of the structure and energetic function of muscle mitochondria stratified based on their association with capillaries in order to evaluate these predictions. Consequently, the aim of this study was to evaluate the impact of capillary and sarcolemmal proximity on mitochondrial structure and energetic function in mouse hindlimb skeletal muscle using 3D ultrastructural and live cell functional imaging approaches.

**Figure 1.**
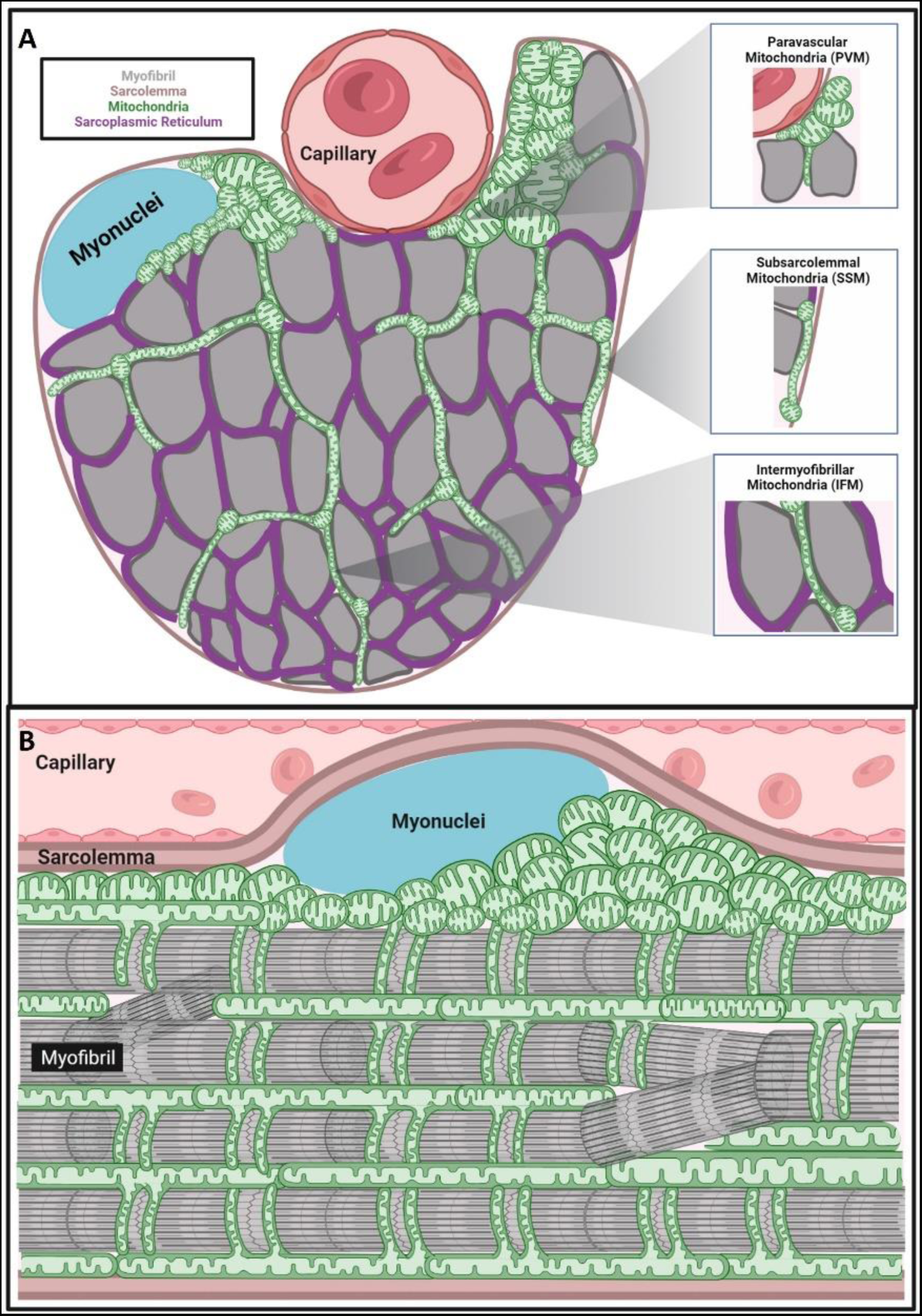
Subcellular region-specific mitochondria. This schematic describes the location and shape differences between the paravascular mitochondria (PVM), interfibrillar mitochondria (IFM), and subsarcolemmal mitochondria (SSM). (A) View of a cross-section of a skeletal muscle cell and where the three spatially specific pools of mitochondria would be located. (B) View of a longitudinal view of a skeletal muscle cell to demonstrate how mitochondria are located within the myofibril matrix.

## Methods

### Ethical approval

Procedures were approved by the National Heart, Lung, and Blood Institute Animal Care and Use Committee. The procedures were performed in accordance with the guidelines described in the Animal Care and Welfare Act (7USC2142§13).

### Animals

C57BL/6N mice, age 2 – 4 months, were purchased from Taconic Biosciences (Rensselaer, NY, USA) and fed ad libitum on a 12-hour light, 12-hour dark cycle at 20 - 26°C. All mice were anesthetized with isoflurane via a nose cone as described for specific experiments below and then euthanized via exsanguination.

### FIB-SEM Sample Preparation

Similar to previous studies (Bleck *et al*., 2018; Kim *et al*., 2021), mice were anesthetized using 2% isofluorane via nose cone. While anesthetized, the skin of the hindlimbs were peeled back and the hindlimb was submerged in a fixative solution (2% glutaraldehyde in 0.1M phosphate buffer, pH 7.2) for 30 minutes. The soleus and gastrocnemius were then excised for an oxidative and glycolytic skeletal muscle, and cut into 1 mm slices. These were placed in a new buffer solution (2.5% glutaraldehyde, 1% formaldehyde, 0.12M sodium cacodylate, pH 7.2) overnight at room temperature. This in vivo immersion fixation technique has been shown to maintain physiological capillary size and shape (Kim *et al*., 2020), mitochondrial diameter (Glancy *et al*., 2015), and myofilament lattice spacing (Katti *et al*., 2022b) in adult mouse skeletal muscles.

These samples were washed five times with three minute durations in a 0.1 M cacodylate buffer at room temperature. Samples were reduced in a 4% osmium solution (3% potassium ferrocyanide, 0.2 M cacodylate, 4% aqueous osmium) for 1 hour on ice, washed 5 times in bi-distilled water, and then incubated again in a thiocarbohydrazide solution for 20 minutes at room temperature. Next, samples were incubated in a 2% osmium solution for 30 minutes on ice, and then washed again with bi-distilled water. The samples incubated overnight in a 1% uranyl acetate solution at 4°C. The next day, samples were washed with bi-distilled water, incubated at 20°C for 20 minutes in a Walton’s lead aspartate solution (0.02 M lead nitrate, 0.03 M aspartic acid, pH 5.5), and washed again five times with bi-distilled water. The samples were dehydrated with decreasing concentration of ethanol (20%, 50%, 70%, 90%, 95%, 100%; 5 minutes each) and then incubated in a 50% Epon solution (50% ethanol) for 4 hours and incubated in 75% Epon resin (25% ethanol) overnight at room temperature. The next day, samples were incubated in fresh 100% Epon resin for 1 hour, 1 hour, and 4 hour incubations at room temperature. Samples were placed onto aluminum Zeiss SEM Mounts after removing excess resin and polymerized in a 60°C oven for 48 hours. After polymerization, stubs were mounted in a Leica UCT Ultramicrotome (Leica Microsystems Inc., USA) and faced with a Trimtool 45 diamond knife (DiATOME, Switzerland) at a feed of 100 nm at a rate of 80 mm/s.

### FIB-SEM Imaging

Images were acquired with a ZEISS Crossbeam 540 using the ZEISS Atlas 5 software (Carl Zeiss Microscopy GmbH, Jena, Germany) and were collected using an In-Column Energy Selective Backscatter (ESB) with filtering grid to reject unwanted secondary electrons as well as backscatter electrons up to a voltage of 1.5kV at the working distance of 5.01 nm. The milling was performed with a FIB operating at 30 kV and 2 – 2.5 nA beam current. The thickness of the FIB slices are 5 – 10 nm. Image stacks were aligned with proprietary algorithm by using the Atlas 5 software (Fibics) and exported as TIFF format for future analysis.

### Segmentation of cellular structures

Segmentation of myosin, mitochondria, lipid droplets, sarcotubular network (SR/T), and z-disks were performed using the Pixel Classification module in the Ilastik software package(Sommer *et al*., 2011). Notably, the sarcotubular network represents the abundant interaction of the sarcoplasmic reticulum (SR) and the t-tubules, resulting in the majority of the t-tubule surface covered by SR(Franzini-Armstrong, 1991). Raw images of skeletal muscle containing a capillary were loaded sequentially as 8 bit TIF images into ImageJ (National Institutes of Health, Bethesda, MD) and exported as a single HDF5 file using the Ilastik plugin before being loaded into Ilastik. Pixel classification was completed by tracing cellular structures in the XY, XZ, and YZ planes of the sample volume. Classification labels were made for the outer mitochondrial membrane, inner mitochondria, myosin, z-disks, lipid droplets, and SRT. After initial training, “Live Update” tool was used to display the results of the segmentations, and iterative corrections were made for erroneous mislabeling errors. Training continued until satisfactory segmentation was observed. The pixel probability files were then exported as 32-bit HDF5 files.

To segment individual mitochondria, the pixel probabilities files were loaded into the MultiCut module in the Ilastik software. The mitochondrial outer membrane label was overlaid onto the raw image files to create super pixels. The super pixels were analyzed to ensure boundaries of the mitochondria were followed. If this was not the case, the threshold was adjusted and new super pixels were generated until satisfaction. The boundaries of the super pixels was then performed by selecting the mitochondria membrane as “red” and the non-mitochondrial boundaries as “green”. The “Live Predict” tool was then used to establish the amount of training needed. Upon appropriate training, accurate predictions were made with the “Live Multicut” tool with the Nifty_FmGreedy solver and 0.3-0.5 beta to generate the individual mitochondrial segmentations. The multicut segmentation was then exported as a 32-bit HDF5 file for analysis.

### Segmentation of mitochondrial pools

It has been proposed that the embedding of capillaries into the muscle fiber, rather than direct interactions with the sarcolemma, creates myofibril voids where large peripheral mitochondrial pools form (Glancy *et al*., 2014; Glancy & Balaban, 2021; Willingham *et al*., 2021). However, mitochondria are not equally dispersed on all sides of the embedded capillary. Often, mitochondria are clustered in pools on either side of the capillary, and these pools thin out at increasing distances away from a capillary. The change in geometry around the capillary therefore results in inconsistent selection of mitochondria when using distance from capillary as a relative marker. To overcome this, the maximum distance of the mitochondrial outer membrane from myosin was the chosen thresholding marker. We reasoned mitochondria closest to myosin were most likely to be a part of the interfibrillar mitochondrial pool, as interfibrillar mitochondria are surrounded on all sides by the contractile apparatus. Paravascular mitochondria, however, are surrounded by the contractile apparatus on one side or not at all. Therefore, we predicted that mitochondria with smaller maximum distances to myosin were most likely to be a part of the interfibrillar mitochondrial pool and the mitochondria with larger maximum distances from myosin to be a part of the paravascular pool. Due to the variability how deep a capillary will embed itself into a muscle cell (Glancy *et al*., 2014), the distance threshold was optimized for each cell.

To perform the paravascular mitochondria selection, HDF5 files for total mitochondria segmentation and myosin segmentation from the completed multicut segmentation in ilastik was imported into Fiji/ImageJ. The HDF5 files were saved as 8-bit TIFF images prior to any additional analysis. A manual threshold of 128 to 255 was selected to make a binary myosin image stack. A distance map of was created from an 8-bit grey myosin image stack using “Distance Transform 3D” in the Process plug-in. A table of the minimum, maximum, and mean myosin distance values was created using “Intensity Measurements 2D/3D” function within the MorphoLibJ plug-in. The myosin distance map was the input and the total mitochondria image stack was selected as the labels. The maximum myosin distance was assigned to the mitochondrial labels using the “Assign Measure to Label” tool in the MorphoLibJ plug-in. An optimal threshold value was then determined for each cell in order to ensure a minimum of 90% paravascular mitochondria were selected and no more than 10% false selection of non-paravascular mitochondria. For the three oxidative cells used in this study, we determined a minimum threshold of 560 nm, 600 nm, and 640 nm to provide optimal paravascular mitochondria selection. Erroneous selections of mitochondria not next to the capillary were individually removed with the “Label Edition” tool of the MorphoLibJ plug-in. The threshold image was transformed into a binary image, which only included the paravascular mitochondria. The “Image Calculator” tool in ImageJ was used to multiply the binary paravascular mitochondria pool and the originally mitochondrial labels resulting in an image of mitochondria labels in the paravascular mitochondria pool which was then saved as 32-bit TIF files. Using this selection method, mitochondria which extend from the paravascular region into the intrafibrillar region (Glancy *et al*., 2015) are included as PVM and not IFM.

We defined subsarcolemmal mitochondria as mitochondria located directly under the sarcolemma but not associated with the paravascular pool. To select these mitochondria, a distance map of the sarcolemma was made with the “Distance Transform 3D” tool in ImageJ. A table of the minimum, maximum, and mean sarcolemma distance values was created using “Intensity Measurements 2D/3D” function within the MorphoLibJ plug-in. The minimum distance from the sarcolemma was assigned to the mitochondrial labels using the “Assign Measure to Label” tool. A threshold of 0 to 40 nm was then applied. To exclude mitochondria also in paravascular mitochondria, the paravascular mitochondria selection was subtracted from the 40 nm thresholded subsarcolemmal mitochondria using the “Image Calculator” tool in ImageJ. The resulting mitochondria are the SSM. The binary subsarcolemmal mitochondria was multiplied by the original mitochondrial labels using the “Image Calculator” and was saved as a 32-bit TIF file.

To create the interfibrillar mitochondria pool, a binary image of 0 and 1 was created for the original mitochondrial labels. Using the image calculator tool, the binary paravascular mitochondria pool and subsarcolemmal pool was subtracted from the original binary mitochondrial labels. This resulted in a binary image of only interfibrillar mitochondria. Using the image calculator tool again, the binary interfibrillar mitochondria was multiplied against the total mitochondrial labels to produce an image of only interfibrillar mitochondria pool mitochondria, which was subsequently saved as a 32-bit TIF file.

### Accuracy and validity of paravascular mitochondria selection

To evaluate the automated selection of paravascular mitochondria using maximal distance from myosin rather than distance from the sarcolemma, we determined the accuracy of paravascular mitochondria selection using each of these measures. The total number of mitochondria in the paravascular pool was determined by identifying mitochondria in the paravascular pool manually. Next, the paravascular mitochondria was selected using the 540 nm, 600 nm, or 640 nm and greater distance from myosin, as described above. Correctly selected mitochondria were identified manually and then divided by the total number of paravascular mitochondria for that cell. This was repeated using a 0 – 40 nm distance from sarcolemma threshold. False positive mitochondria were defined as mitochondria selected by the threshold which were not part of the paravascular mitochondria pool. The total number of incorrectly identified mitochondria was divided by the total number of mitochondria selected by each method.

### Individual mitochondrial analysis

Each 32-bit mitochondrial pool TIF file was loaded into ImageJ. Measures of mitochondrial volume, surface area, sphericity, elongation (largest radius/middle radius), and mitochondrial length were measured using the 3D Geometrical Measure tool within the 3D Manager in the 3D Suite plug in. Individual mitochondrial surface area to volume ratio was calculated by dividing surface area and volume values. Individual mitochondrial aspect ratio was calculated by multiplying elongation and flatness values. Mitochondrial length was determined using the “Geodesic Diameter 3D” tool in the MorpholoLibJ plug-in.

### Mitochondria to organelle interactions

To determine the interaction of the mitochondria with other organelles and cellular structures, a mask of the outer mitochondria membrane was created using “Label Boundaries” in the MorphoLibJ PlugIn in each of the 32-bit mitochondrial segmentations. The new outer mitochondria membrane image sequence was divided by itself using the Image Calculator with a 32-bit (float) result to yield values of 1.0 for all outer mitochondrial membrane pixels and NA for all others. The segmented organelles and cellular structures were made into 8-bit gray valued images. Using the “Distance Transform 3D” tool, distance maps from each of the binarized cellular structures were made. The “Intensity Measurements 2D/3D” was used to calculate the minimum distance from the organelle/cell structure with the distance map as the input and the mitochondrial labels as the labels. The mitochondrial labels image was selected, and then assigned new minimum distance labels using the “Assign Labels” tool in the MophoLibJ plugin. A threshold of 40 nm or less was used to determine contact sites with lipid droplets, sarcoplasmic reticulum, and sarcolemma, and the proximity to myosin. The threshold images were multiplied by the original mitochondrial label boundary image and then subtracted from the 32-bit mitochondria segmentation images using the Image Calculator tool. The 3D Geometrical Measure tool in the ROIManager 3D plugin was used to determine the resulted mitochondrial pixel volumes. The percentage of mitochondrial surface area in contact with a given cellular structure was calculated from the difference in pixels between the interactive subtracted pixel volumes and the original mitochondrial pixel volumes.

### Electroporation of Tq-FLITS Plasmid to the Flexor Digitorum Brevis

Electroporation was performed similarly to described methods(Hain & Waning, 2022). Briefly, animals were anesthetized with 5% isoflurane and placed in a prone position. A total of 30 μL of 2mg/mL hyaluronidase was injected below the skin of the foot and above the flexor digitorum brevis muscle. The animal then recovered from anesthesia for 1 hour. Again, the animal was anesthetized with 5% isoflurane and placed in a prone position. Another injection of 30 μL of the mitochondrially targeted Tq-FLITS plasmid (mitoTq-FLITS) (van der Linden *et al*., 2021), generously donated by Elizabeth Murphy’s Laboratory, NHLBI, NIH, was injected below the skin of the foot.

After ten minutes, electrodes were inserted into the FDB to create an electric field through the muscle. Electrical stimulation was applied for 20 seconds with each electric pulse occurring every second at 100 volts of electricity. A minimum of two weeks were allowed for uptake and production of the mitoTq-FLITS protein prior to flexor digitorum fiber isolation and imaging.

### Flexor Digitorum Brevis (FDB) Fiber Isolation & Imaging

Muscle fiber isolation and imaging was performed similarly to previously established methods(Park *et al*., 2014; Liu *et al*., 2020). Animals were anesthetized with 5% isoflurane and euthanized. Immediately after euthanasia, the FDB muscles of mice were dissected and places into warm Tyrodes buffer (pH 7.4, 10 mM HEPES, 137 mM sodium chloride, 4.5 mM potassium chloride, 0.5 mM magnesium sulfate, 0.5 mM potassium phosphate, 10 mM glucose, and 1.8 mM calcium chloride) and 3 mg/ml collagenase solution. The muscles were agitated in a water bath at 37°C for 75 minutes. Individual muscle fibers were released by gentle trituration of the FDB muscle in new Tyrodes solution(Park *et al*., 2014). Once cells were isolated, the solution was set aside to allow cells to settle to the bottom of the tube. The remaining Tyrodes solution was syphoned off and cells were incubated in 5 nM TMRM and 200 nM MitoTracker Green dye at 37°C for 20 minutes prior to imaging.

Isolated muscle cells were imaged with a Leica upright SP8 microscope with a Nikon 25x (1.1 NA) water-immersion objective. Confocal sequential line scanning and internal HyD detectors were used during image capture. MitoTracker Green and TMRM were imaged with 488 nm and 552 nm excitation and 500 – 550 nm and 590 – 650 nm emission, respectively. Pixel size was 0.171 μm and a line average of 256 was used for image capturing.

Image analysis of the TMRM and MitoTracker Green was performed in ImageJ. A mask of mitochondrial content was created by using the Threshold tool on the MitoTracker Green image to remove any background pixels. The mask was then applied to the TMRM image to normalize the mitochondrial membrane potential to the pixels contained within the mitochondrial content mask. Using the Image Calculator in ImageJ, the mitochondrial mask was divided by itself to obtain 1 and NA values. The resultant image was then multiplied by the TMRM and MitoTracker Green image using the Image Calculator to obtain pixel normalized images. Using the free hand selection tool, paravascular mitochondria ROI was selected. The rectangle tool was then used to obtain a paired measurement of IMF immediately adjacent to the paravascular mitochondria ROI, as a way to account of any optical differences that may be seen within one fiber. The subsarcolemmal mitochondria were selected on the same optical plane as the paravascular mitochondria and interfibrillar mitochondria. Mitochondria from the edge of the cell to 2 μm interior of the cell were selected as part of the subsarcolemmal mitochondria to be sure all subsarcolemmal mitochondria were included. The mean gray pixel values of each ROI were then used to determine pixel intensity.

Imaging of the mitoTq-FLITS probe in isolated skeletal muscle fibers was performed using the fluorescent lifetime imaging (FLIM) module on a Leica SP8 STED 3X/Confocal Microscope. Fibers were excited at 480 nm and collected at emission wavelengths of 500 – 670 nm. To obtain enough photon counts, a 35 line average was used. Lifetime analysis was performed with a two component exponential model. The amplitude weighted mean lifetime was taken for each hand drawn ROI for the PVM, IFM, and SSM.

### NADH Redox and Flux

Mitochondrial energetic flux was measured using the MitoRACE method (Willingham *et al*., 2019). Briefly, FDB fibers were isolated as described above. The isolated fibers were then plated on Cell-Tak coated dishes. Optimal adhesion was achieved by allowing the cells to settle for upwards of 60 minutes before exposing the cells to perfusion.

Imaging cells was performed using a Leica upright SP8 microscope. Multiphoton laser-scanning excitation was then used to collect the autofluorescence of NADH. A micro perfusion system (MultiChannel Systems, Reutlingen, Germany) was then used to perfuse Tyrodes buffer through the coated cells at 7.8 mL/min. After 30 seconds of base-line reading, 5 mM sodium cyanide was added to solution to inhibit oxidative phosphorylation. The immediate rise in NADH autofluorescence was then captured.

NADH fluorescence was excited at 750 nm and emission was detected using a Leica HyD detector at 414 – 538 nm. Rapid volumetric imaging of 32 μm in depth was also employed to obtain full cell-volume measurement(Bakalar *et al*., 2012). ROIs for PVM, SSM, and IFM were drawn as described above for the TMRM and mitoTq-FLITS analysis. Importantly, we previously showed (Willingham *et al*., 2019) that fully oxidized NADH results in zero signal in our system and that both mitochondrial NADPH and cytosolic NAD(P)H contribute less than 5% of the cellular fluorescent NAD(P)H signal in adult mouse skeletal muscle.

### Statistical Analysis

Linear regression analysis was used to assess individual mitochondrial morphology and connectomics measures of the three region specific mitochondria pools. Linear regression analysis was also used to determine whole cell mitochondria interaction with organelles.

Mitochondrial function was assessed using multilevel models with TMRM fluorescence, NADH redox, NADH flux, and mitoTq-FLITS Lifetime as dependent variables. Typically, pixel intensity measures for TMRM would be divided by MTG to get a ratio of fluorescence per unit of mitochondrial content. However, representations of these data through ratios require certain statistical assumptions to be met(Goran *et al*., 1995; Curran-Everett, 2013). After dividing TMRM by MTG to get a ratio of TMRM/MTG, the new, ratio variable still correlated with MTG (r= −0.22, p=0.006) meaning that dividing by MTG did not get rid of mitochondrial content’s influence on TMRM fluorescence. Further, when regressing TMRM on MTG, there is a non-zero intercept (β_0_=2.76, p<0.001), which was a substantially better predictor of TMRM than a model with a zero-fixed intercept (p<0.001). Therefore, neither of the primary assumptions to create a ratio variable were met, so we analyzed the data using the more appropriate, model comparison approach(Goran *et al*., 1995; Curran-Everett, 2013).

To calculate differences in TMRM fluorescence between mitochondrial pools, intercepts were allowed to vary across fiber. In each regression model, MTG was added into the model prior to adding variables of interest to account for mitochondrial content, which is superior to the commonly used normalization method(Goran *et al*., 1995; Curran-Everett, 2013). A categorical mouse variable was then entered to account for naturally occurring biological differences between mice. Type of mitochondrial pool (SSM, IFM, or PVM) were added to the model to determine their effect of functional measure above and beyond mitochondrial content and biological variability.

For both NADH redox, NADH flux, and mitochondrial calcium measurements, intercepts were allowed to vary across fiber. Mouse was entered into the regression equation first to account for biological differences. Mitochondrial pool were entered as independent variables to analyze their effect after accounting for biological variability. All statistical analyses were performed in *R Studio* (Version 4.1.1).

## Results

### Accuracy of mitochondrial subcellular region segmentation

To determine the optimal method of selecting paravascular mitochondria (PVM) in an automated fashion within our FIB-SEM datasets, we evaluated the regional assignment of every mitochondrion based on either their minimum distance from the sarcolemma or the maximal distance from myosin (Figure 2A – 2D). Maximal distance from myosin resulted in a 93.85±1.86% (mean ± SD) correct selection of PVM whereas minimum distance from the sarcolemma resulted in 73.57±9.02% correct selection of PVM (Figure 2E). Moreover, using the minimum distance from the sarcolemma resulted in a false positive selection of non-PVM of 31.97±12.53% compared to a 5.03±4.42% false positive rate when using maximal distance from myosin (Figure 2F). Thus, we determined that maximal distance from myosin was the superior automated method to select paravascular mitochondria.

**Figure 2.**
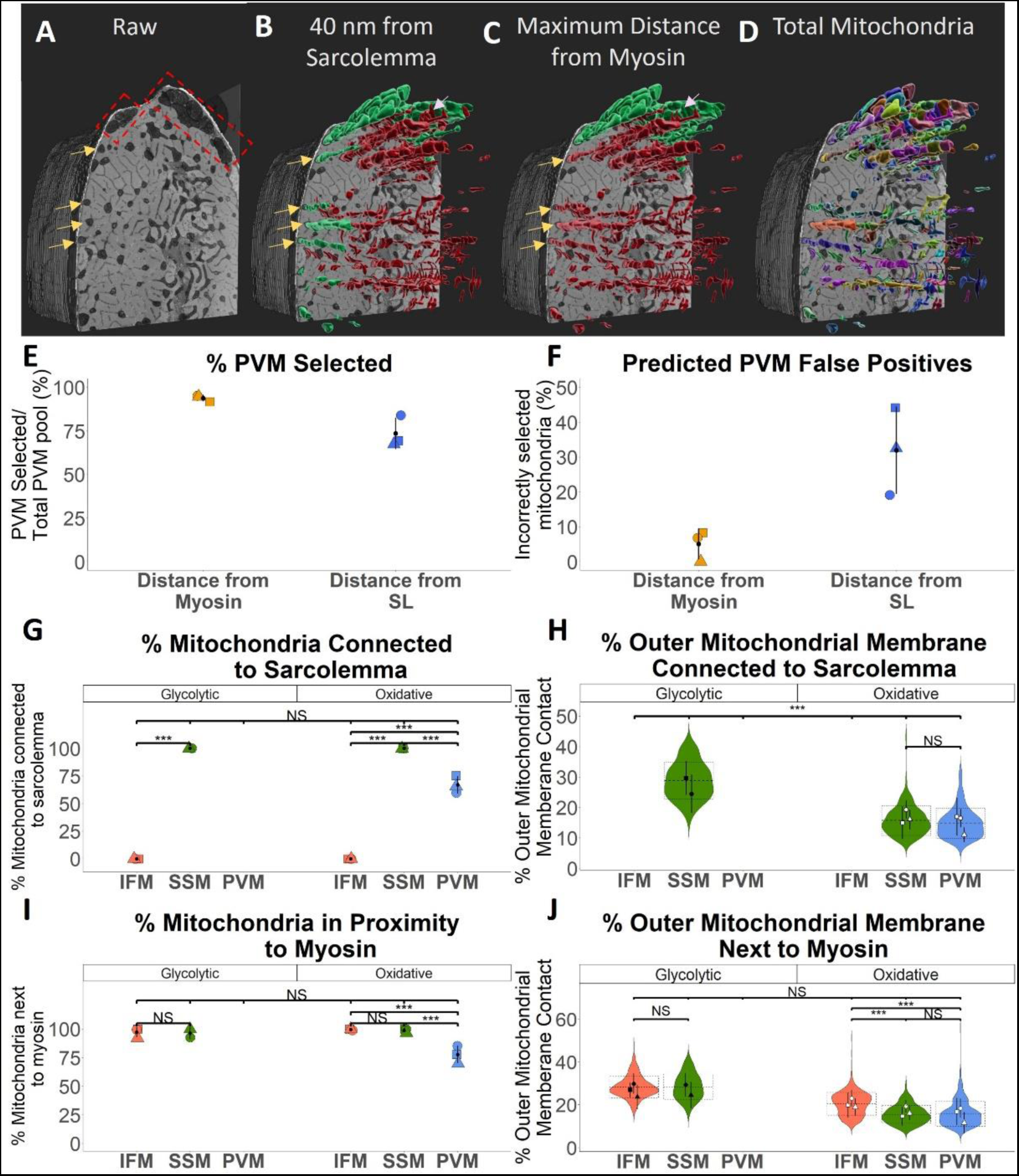
Accuracy and validation of separating interfibrillar, subsarcolemmal, and paravascular mitochondria. (A-D) Representative images of the different methods of separating the pools of mitochondria. In the red box in (A), the identified pools of mitochondria are seen. The yellow arrow identifies the mitochondria under the sarcolemma that are not associated with the rest of the mitochondria pool. The white arrow identifies mitochondria selected by the maximum distance from myosin but not the minimum distance from the sarcolemma. Mitochondria selected by each method are observed in green and the mitochondria not selected are shown in red. Using the minimum distance from the sarcolemma, both the pooled mitochondria and the mitochondria identified by the yellow arrow are selected. Using the maximum distance from myosin, only the pooled mitochondria are selected. (E) The accuracy measurements when selecting the paravascular mitochondria using maximal distance from myosin and minimum distance from the sarcolemma (SL). (F) The false positive percentage when selecting the paravascular mitochondria using maximal distance from myosin and minimum distance from the SL. (G) The total amount of mitochondria (n=4,000) in contact with the SL in each mitochondria pool for glycolytic (n=3) and oxidative cells (n=3). (H) The percentage of the outer mitochondrial membrane in contact with the SL in each mitochondrial pool for glycolytic and oxidative cells. (I) The total percentage of mitochondria near myosin in each mitochondrial pool for glycolytic and oxidative cells. (J) The percentage of the outer mitochondrial membrane next to myosin in each mitochondrial pool for glycolytic and oxidative cells.

To further validate our PVM selection method, we determined what percentage of mitochondria were in contact with the sarcolemma (Figure 2G). For both glycolytic and oxidative cells, IFM had 0% contact with the sarcolemma, SSM had 100% contact with the sarcolemma, and PVM had 66.20% contact with the sarcolemma. This is to be expected, as none of the IFM should be in contact with the sarcolemma, all of the SSM should be in contact with the sarcolemma, and only some of the PVM will be in contact with the sarcolemma. Of the mitochondria that were in contact with the sarcolemma, the relative amount of outer mitochondrial membrane making sarcolemmal contact was not different between SSM and PVM in oxidative cells (Figure 2H).

Lastly, we confirmed that IFM and SSM were in proximity to myosin whereas only some of the PVM would be in proximity to myosin. For IFM and SSM in both glycolytic (98.57±11.88% and 93.75±24.36%) and oxidative cells (99.87±6.42% and 99.21±8.87%), nearly all mitochondria were found to be in proximity of myosin. Moreover, only 78.09±41.41% of PVM were in proximity to myosin (Figure 2I). Again these results were as expected, as IFM and SSM will be located directly next to myosin by definition, whereas only the deeper mitochondria of the PVM will be located next to myosin with the remaining mitochondria located closer to the sarcolemma with no contact to myosin. Finally, in glycolytic cells, the amount of outer mitochondrial membrane next to myosin was no different between IFM and SSM. In oxidative cells, IFM had more outer mitochondrial membrane in proximity to myosin compared to both SSM and PVM, with no difference observed between SSM and PVM (Figure 2J). Thus, the above data confirm that our automated segmentation routine accurately selected PVM, SSM, and IFM.

### Individual mitochondrial morphology comparison

To evaluate mitochondrial structural differences across subcellular regions, individual mitochondrial morphology was compared for IFM, SSM, and PVM. In oxidative cells, both PVM (0.60±0.69 μm^3^) and SSM (0.59±0.57 μm^3^) were larger than IFM (0.31±0.42 μm^3^, p<0.001; Figure 3E). IFM had the largest surface area to volume ratio (13.06±3.46) with no differences observed between the SSM (9.76±2.71) and PVM (10.40±4.3, p<0.001; Figure 3F). Sphericity was highest in the PVM (0.42±0.18), with no differences observed between the SSM (0.38±0.16) and IFM (0.36±0.18; Figure 3G). Aspect ratio was lowest in the PVM (2.70±1.16) and similar between IFM (3.48±2.32) and SSM (3.44±2.54, p<0.001; Figure 3H). Lastly, mitochondrial elongation and length was highest in the SSM (2.27±1.62 and 1.74±1.4) compared to both IFM (2.21±1.38 and 1.60±1.17) and PVM (1.83±0.81 and 1.41±0.78, p<0.05 and p<.001; Figure 3I and Figure 3J). Together, these data suggest that PVM are more compact than either IFM or SSM and may be structurally designed for higher internal capacity for energy production (Glancy *et al*., 2020).

**Figure 3.**
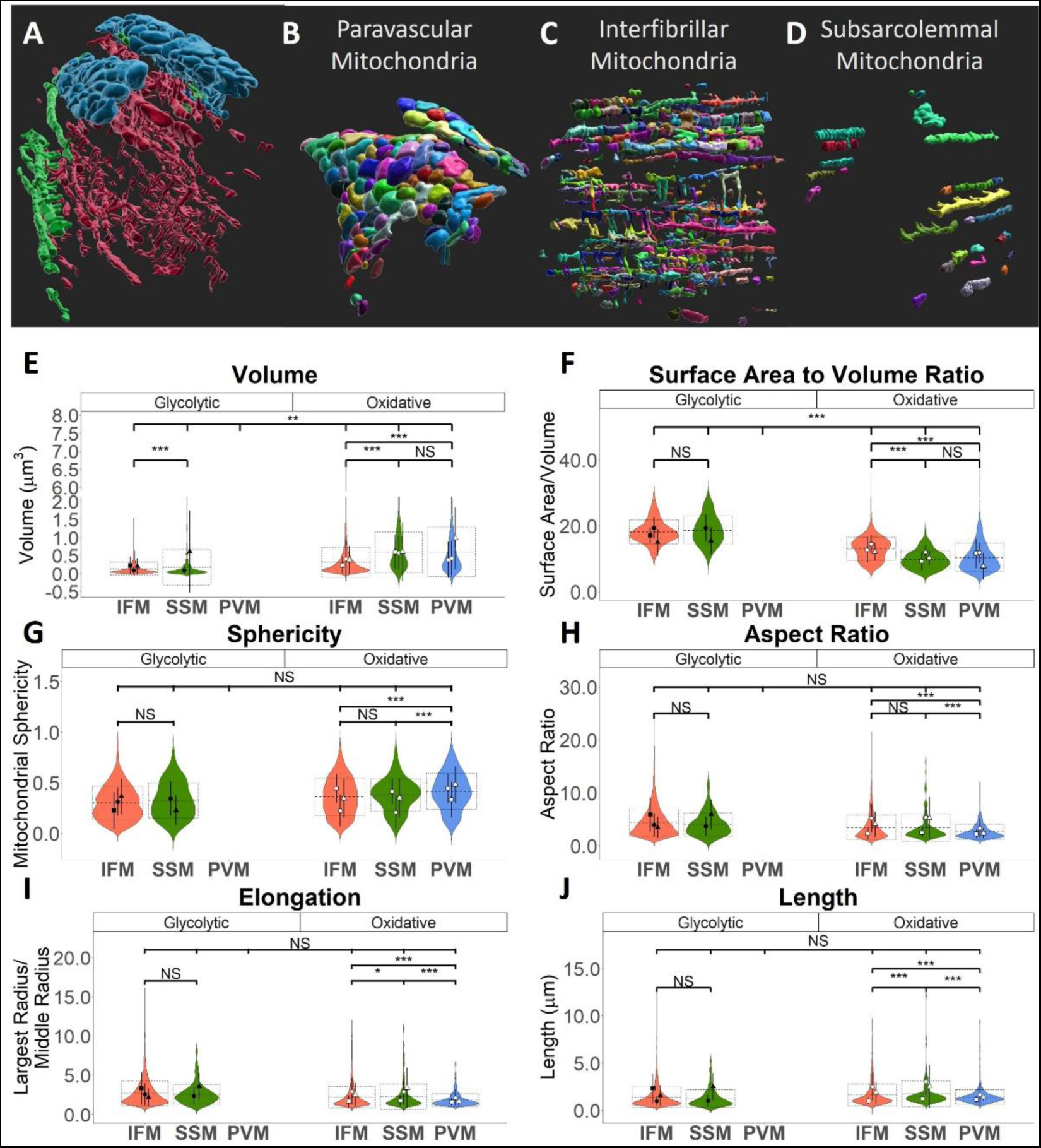
Individual mitochondrial morphology of the IFM, SSM, and PVM in glycolytic and oxidative skeletal muscle cells. (A-D) Representative images of the mitochondrial pools together within one muscle cell, and individually separated. (E) Individual mitochondrial volume, (F) surface area to volume ratio, (G) sphericity, (H) aspect ratio, (I) elongation, and (H) length measured and compared between the mitochondrial pools and fiber type. Mean and standard deviation of each cell are shown as black filled and white filled shapes for glycolytic and oxidative cells, respectively. A square symbol is for cell 1, a circle is for cell 2, and a triangle is for cell 3 of each fiber type. Group mean and standard deviations are shown with a black dot and solid line. * represents p < 0.05. ** represents p < 0.01. *** represents p < 0.001.

To further evaluate the influence of the sarcolemma on mitochondrial structure without the influence of embedded capillaries, we also assessed glycolytic fibers which have little or no PVM due to a lack of embedded capillaries (Glancy *et al*., 2014). Similar to oxidative cells, SSM were slightly larger (0.18±0.49 μm^3^) compared to IFM (0.14±0.41 μm^3^; p<0.001) in glycolytic cells. Conversely, SSM and IFM had similar surface area to volume ratio, sphericity, aspect ratio, elongation, and mitochondrial length (p>0.05) in glycolytic cells. These data suggest that the sarcolemma alone has minimal influence on mitochondrial shape.

### Mitochondrial connectivity comparison

To investigate how mitochondrial subcellular location impacts structural capacity for interorganelle communication and exchange, we evaluated the connectivity among mitochondria, lipid droplets, and the SR in each subcellular region (Figures 4-6). No lipid droplets were found in the glycolytic fibers, precluding mitochondria-lipid droplet analyses in these cells. The percentage of mitochondria in contact with lipid droplets in oxidative cells was not significantly different among subcellular pools (Figure 4C). However, of the mitochondria in contact with lipid droplets, the IFM had a higher percentage of outer mitochondrial membrane in contact with lipid droplets (18.91±5.34%) compared to either the SSM (14.69±4.20%) and PVM (15.18±5.06%, p<0.001; Figure 4D). No differences in mitochondria-lipid droplet connectivity were detected between SSM and PVM.

**Figure 4.**
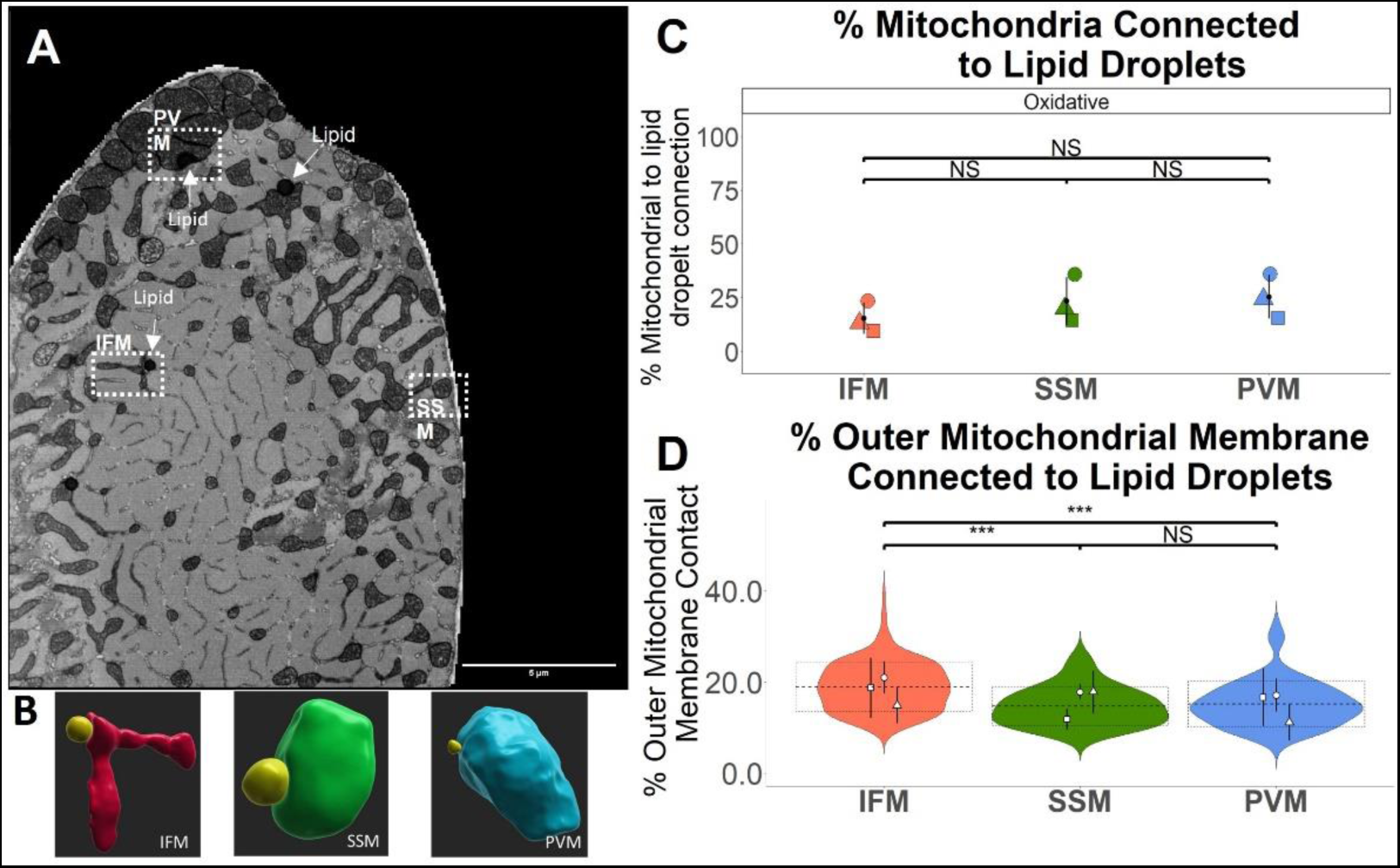
Mitochondria-to-lipid droplet interaction. (A) Representative image of an electron microscopy slice of a skeletal muscle cell showing the location of paravascular mitochondria, subsarcolemmal mitochondria, interfibrillar mitochondria, and a lipid droplet. Scale bar is 5 μm. (B) Representative renderings of mitochondria from each pool in contact with a lipid droplet (red = interfibrillar mitochondria, green = subsarcolemmal mitochondria, blue = paravascular mitochondria, yellow = lipid droplet). (C) The percent of mitochondria located 40 nm or less from a lipid droplet. (C) The percentage of outer mitochondrial membrane in contact with the surface of lipid droplets A square symbol is for cell 1, a circle is for cell 2, and a triangle is for cell 3 of each fiber type. A black dot and solid line show group means and standard deviations. * represents p < 0.05. *** represents p < 0.001. Mitochondria and lipid droplet interaction is only shown in oxidative cells.

Consistent with our previous results (Bleck *et al*., 2018; Kim *et al*., 2021), nearly every muscle mitochondrion was in contact with the SR regardless of cell type or subcellular region (Figure 5C) further indicating the critical importance of mitochondrial-SR interactions in skeletal muscle. Also similar to our earlier work (Bleck *et al*., 2018), glycolytic cells had greater outer mitochondrial membrane contact with the SR compared to oxidative cells (p<0.001; Figure 5D). While there was no difference in relative mitochondrial outer membrane-SR contact between IFM and SSM in glycolytic cells (Figure 5D), IFM in oxidative cells had more outer mitochondrial membrane contact with the SR (20.51±5.30%) compared to both SSM (16.18±6.65%) and PVM (15.49±4.21%, p < 0.001; Figure 5D). No differences in mitochondria-SR connectivity were detected between SSM and PVM in oxidative cells (Figure 5D).

**Figure 5.**
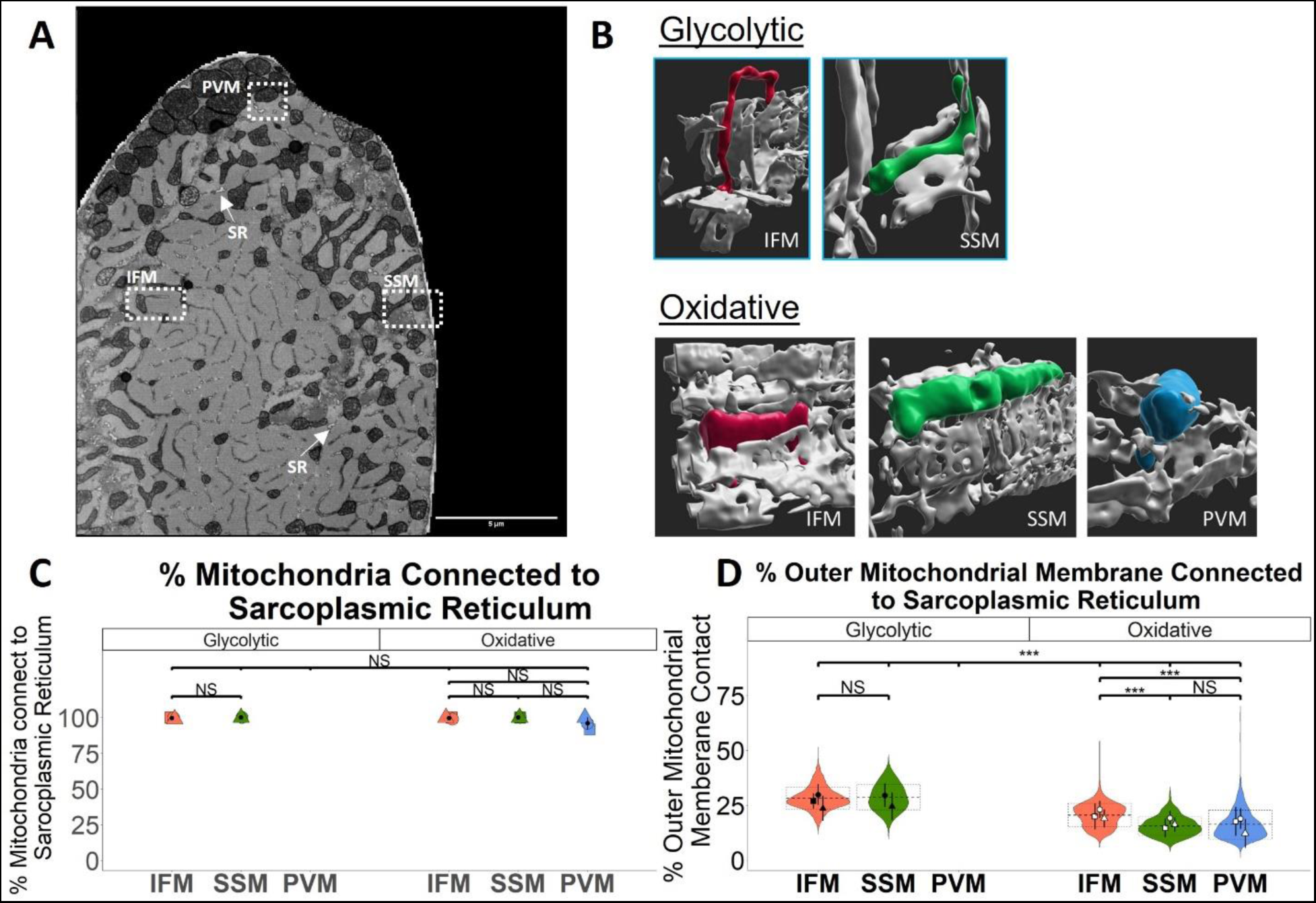
Mitochondria-to-sarcoplasmic reticulum interaction. (A) Representative image of an electron microscopy slice of a skeletal muscle cell showing the location of paravascular mitochondria, subsarcolemmal mitochondria, interfibrillar mitochondria, and sarcoplasmic reticulum. Scale bar is 5 μm. (B) Representative renderings of mitochondria from each pool in contact with the sarcoplasmic reticulum (red = interfibrillar mitochondria, green = subsarcolemmal mitochondria, blue = paravascular mitochondria, white = lipid droplet). (B) The percent of mitochondria located 40 nm or less from the sarcoplasmic reticulum. © The percentage of the outer mitochondrial membrane in contact with the surface of the sarcoplasmic reticulum. A square symbol is for cell 1, a circle is for cell 2, and a triangle is for cell 3 of each fiber type. A black dot and solid line show group means and standard deviations. * represents p < 0.05. *** represents p < 0.001. Mitochondria and lipid droplet interaction is only shown in oxidative cells.

Mitochondria were more likely to form intermitochondrial junctions (IMJs) (Bakeeva *et al*., 1981; Glancy *et al*., 2015; Picard *et al*., 2015) with other mitochondria in oxidative skeletal muscle cells than in glycolytic cells regardless of subcellular region (p<0.05; Figure 6C). Of the IMJ forming mitochondria in oxidative cells, PVM had the highest outer mitochondrial membrane interaction with other mitochondria (33.59±18.76%) followed by SSM (28.06±18.72%) and then IFM (16.04±15.07%; p<0.001; Figure 6D). There was no difference in mitochondria to mitochondria interactions between IFM and SSM in glycolytic cells (p>0.05; Figure 6D).

**Figure 6.**
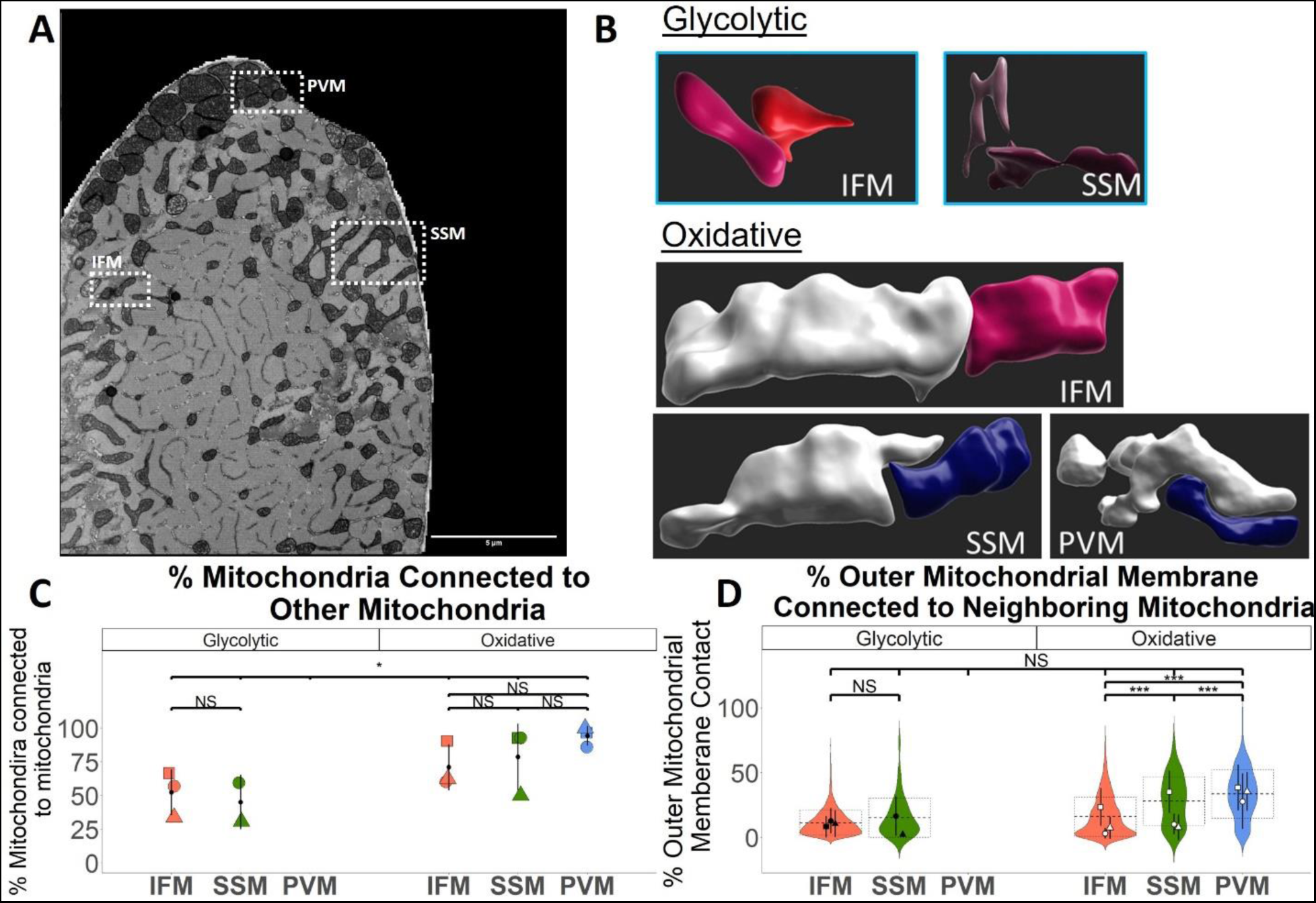
Mitochondria-to-mitochondria interaction. (A) Representative image of an electron microscopy slice of a skeletal muscle cell showing the location of paravascular mitochondria, subsarcolemmal mitochondria, and interfibrillar mitochondria. Scale bar is 5 μm. (B) Representative renderings of mitochondria from each pool in contact with neighboring mitochondria (different colors = different mitochondria). (B) The percent of mitochondria located 40 nm or less from another mitochondrion. (C) The percentage of the outer mitochondrial membrane in contact with the surface of another mitochondrion. A square symbol is for cell 1, a circle is for cell 2, and a triangle is for cell 3 of each fiber type. A black dot and solid line show group means and standard deviations. * represents p < 0.05. *** represents p < 0.001. Mitochondria and lipid droplet interaction is only shown in oxidative cells.

### Functional comparison of the mitochondrial pools

To determine the functional capacity of muscle mitochondria in each subcellular region (Figure 7), we first assessed the basal rate of energy conversion using our recently developed spatially resolved mitochondrial (NADH) flux assay, mitoRACE (Willingham *et al*., 2019). Thus, we were able to evaluate mitochondrial energetic function concurrently in each subcellular region of live individual mouse FDB fibers. The rate of NADH production was highest in PVM (1.63±0.49% NADH reduced per second) followed by SSM (1.45±0.37% NADH reduced per second) and then IFM (1.22±0.36% NADH reduced per second, p<0.05; Figure 7E). To determine if increased driving forces resulted in the greater PVM flux, we also assessed the basal NADH redox state acquired simultaneously with the flux measurement. However, PVM had the most oxidized pool of NADH (50.86±9.35%) compared to both IFM (63.54±5.20%) and SSM (61.04±5.40%, p<0.001; Figure 7F) indicating that the greater PVM flux was achieved with the lowest input driving force in the electron transport chain.

**Figure 7.**
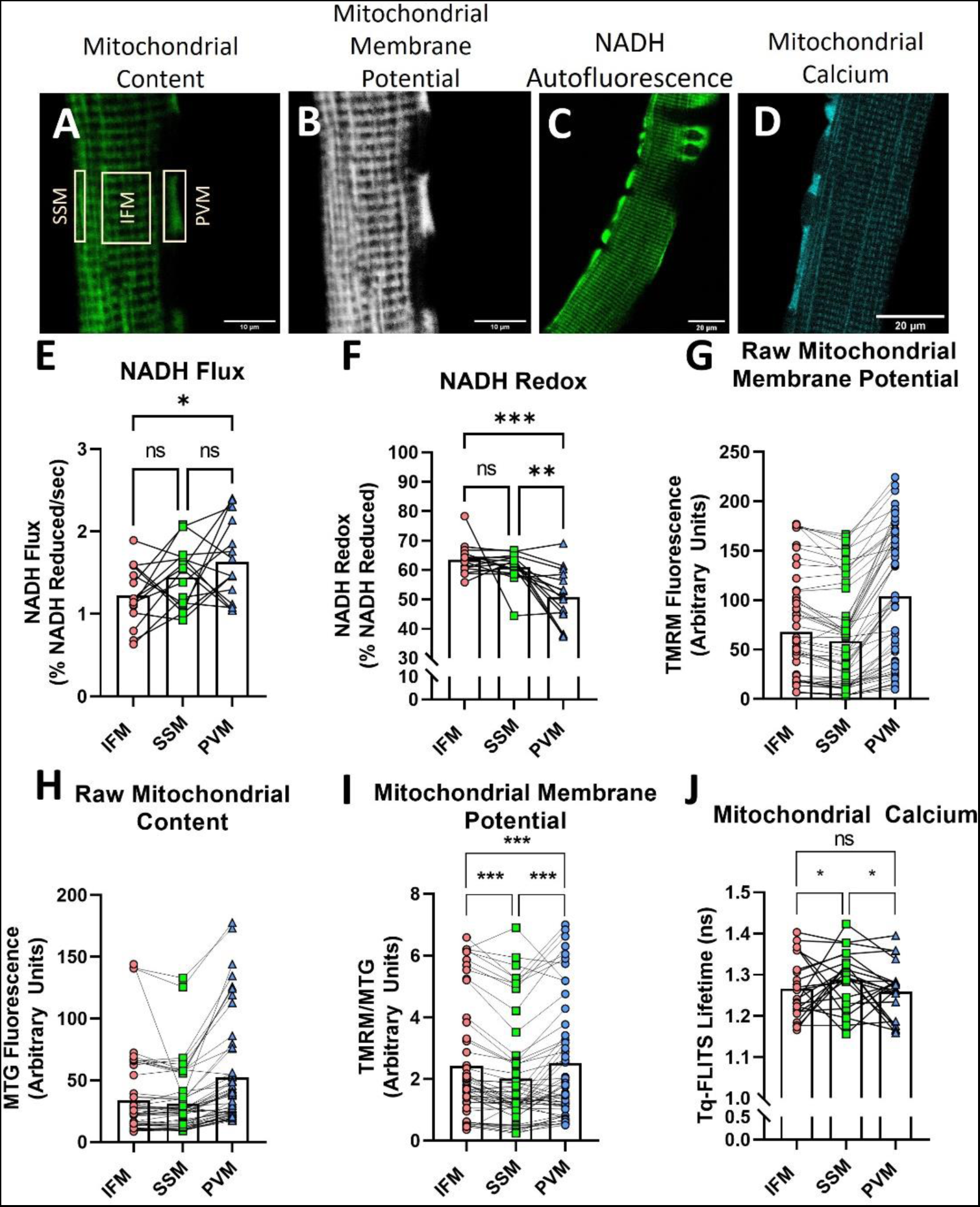
Functional measures of subcellular region-specific pools of mitochondrial. Representative images of mitochondrial content (A), mitochondria membrane potential (B), NADH autofluorescence (C), and mitochondrial calcium levels (D). Panel A also shows the specific regions of interest used to define PVM, IFM, and SSM. (E) NADH flux as measured by NADH autofluorescence pre and post cyanide infusion. (F) NADH redox state pre cyanide addition. (G) Raw mean grey values for mitochondrial membrane potential. (H) Raw mean grey values for mitochondrial content. (I) Mitochondrial membrane potential accounting for mitochondrial content. (J) Tq-FLITS lifetime values were measured for mitochondrial calcium levels. The red circle values represent IFM. The green triangles represent SSM. The blue squares represent PVM. Bars represent the group mean. * represents p<0.05. ** represents p<0.001. *** represents p<0.0001.

To determine if the increased PVM flux was due to a decrease in electron transport chain backpressure, we measured mitochondrial membrane potential using the mitochondrial voltage sensitive fluorophore, TMRM (Figure 7G). Since TMRM fluorescent intensity results from a combination of the voltage across the inner membrane and the content of mitochondria for any given pixel in the image, we concurrently assessed mitochondrial content with the membrane potential-insensitive dye, MitoTracker Green (Pendergrass *et al*., 2004) (Figure 7H). When accounting for mitochondrial content, mitochondrial membrane potential was higher in the PVM compared to SSM and IFM (p<0.001; Figure 7I). Thus, the increased PVM flux occurred in the presence of a lower input driving force and a higher backpressure on the electron transport chain. These data suggest that the effective activity, or conductance (Glancy *et al*., 2013; Fisher-Wellman *et al*., 2018), of the electron transport chain is higher in PVM than in SSM or IFM.

Calcium has long been known to activate mitochondrial metabolism (McCormack *et al*., 1990), including increasing the conductance of the electron transport chain in isolated skeletal muscle mitochondria (Glancy *et al*., 2013). Therefore, we hypothesized that increased calcium levels may explain the greater PVM flux. Basal mitochondrial calcium levels were measured using a pH-insensitive calcium probe, Tq-FLITS (van der Linden *et al*., 2021), targeted to mitochondria. The mitoTq-FLITS probe evaluates calcium levels based on the lifetime of its fluorescence decay, such that lower lifetimes are indicative of lower calcium levels and higher lifetime values are indicative of higher calcium levels. SSM had significantly higher lifetime (1.29±0.07 ns) compared to IFM (1.27±0.07 ns) and PVM (1.26±0.07 ns; p<0.05; **Figure 7J**) indicating greater calcium in SSM. While the lifetime differences between the groups seem to be statistically significant, we consider this difference to be too modest to exclude small systematic errors in lifetime. Moreover, since the measured lifetimes suggest that mitochondrial calcium is lowest in PVM, these data suggest that the increased electron transport chain activation in PVM is not due to mitochondrial calcium.

## Discussion

Skeletal muscle mitochondria have been typically classified as either subsarcolemmal (SSM) or interfibrillar (IFM) based on their respective location within 2D images of skeletal muscle cells. Here, by combining 3D ultrastructural analyses with subcellular functional assessments in live muscle fibers, we demonstrate that mitochondria associated with capillaries have structural and functional characteristics distinct from mitochondria associated with the sarcolemma or myofibrils. While the more voluminous, spherical mitochondria located near the periphery of the muscle cell have been previously classified as subsarcolemmal (Bubenzer, 1966a; Hoppeler *et al*., 1973; Bakeeva *et al*., 1978a; Kirkwood *et al*., 1986b; Picard *et al*., 2012), we find that these large, compact mitochondria are actually those which are associated with capillaries (paravascular) rather than mitochondria directly in contact with the sarcolemma. Moreover, as predicted based on their large volume and compact shape (Glancy *et al*., 2020), we find that mitochondria located lateral to capillaries (PVM) have greater activation of the electron transport chain compared to mitochondria within the same cell and adjacent to the sarcolemma (SSM) or myofibrils (IFM).

Consistent with previous reports (Bleck *et al*., 2018), nearly all skeletal muscle mitochondria (>97%) were in direct contact with the SR/T. However, IFM had a greater proportion of the mitochondrial outer membrane in contact with the SR/T. These data are consistent with previous reports of high IFM to SR interaction in healthy skeletal muscle fibers (Eisner *et al*., 2014; Tubbs *et al*., 2018), and the increase in SR interactions may allow for more sensitive local calcium regulation of ATP production at the location where ATP demand is highest (Eisner *et al*., 2014; Tubbs *et al*., 2018; Willingham *et al*., 2021). Additionally, IFM had higher outer mitochondrial membrane interaction with lipid droplets. Lipid droplets preferentially localize to the interior region of skeletal muscle cells(Nielsen *et al*., 2010; de Almeida *et al*., 2023), supporting the current observation of higher IFM to lipid droplet interaction. Mitochondria in contact with lipid droplets have larger volumes which may facilitate an increase in ATP production compared to mitochondria not directly attached to lipid droplets (Bleck *et al*., 2018). It would be beneficial for a fuel rich density to be closest to myosin, as this provides the shortest diffusion distance for ATP where energy demand is greatest. Interestingly, the highest mitochondria-to-mitochondria connection was observed in the PVM. Cardiac tissue also has the largest mitochondrial volume and intermitochondrial junctions compared to oxidative or glycolytic skeletal muscle (Bleck *et al*., 2018). The similarities between mitochondria from cardiac tissue and PVM from oxidative skeletal muscle suggest a high intrinsic energetic characteristic of PVM compared to IFM. Furthermore, the increased connection in the PVM may allow for the transfer of the mitochondrial membrane potential to be propagated into the interfibrillar mitochondria (Rothstein *et al*., 2005; Glancy *et al*., 2015; Bleck *et al*., 2018; Willingham *et al*., 2021). Taken together, these data suggest IFM are optimally shaped to facilitate SR/T and fuel deposit interactions to regulate and maintain ATP production whereas PVM are structured to build and distribute the proton motivate force needed to drive oxidative phosphorylation, consistent with the relatively higher abundance of cytochrome oxidase (Complex IV) content in PVM compared to IFM (Glancy *et al*., 2015).

Much of our current understanding of functional differences among mitochondria from different striated muscle subcellular regions originates from studies on mitochondria removed from their cellular environments (Palmer *et al*., 1977; Adhihetty *et al*., 2005; Koves *et al*., 2005; Ferreira *et al*., 2010; Kavazis *et al*., 2017; Lai *et al*., 2019). However, the mitochondrial isolation process necessarily separates mitochondria from their networks and results in spherical mitochondrial shapes, both of which may alter the function of mitochondria compared to their native environment (Picard *et al*., 2010). Here, we overcome these limitations by utilizing recently developed methods for assessing mitochondrial function with spatial resolution in live, intact muscle cells (Willingham *et al*., 2019). Using two-photon imaging, we observed PVM to have the highest basal activity, as measured by NADH flux. However, this greater activity occurred despite the NADH redox state being more oxidized and the mitochondrial membrane potential being greater in the PVM compared to SSM and IFM. Thus, PVM achieved the highest flux rates with the lowest energetic driving force across the electron transport chain indicating a greater conductance along this portion of the energy conversion pathway. These data are consistent with previous studies showing a more oxidized redox state of PVM compared to IFM in intact skeletal muscle cells (Kuznetsov *et al*., 2006; Schroeder *et al*., 2010; Willingham *et al*., 2019). However, we previously reported no difference in basal NADH flux between PVM and IFM (Willingham *et al*., 2019). This discrepancy may be due to the current study using isolated cells rather than an *in vivo* model, the relatively higher SSM rates being grouped together with IFM previously, or the larger number of replicates performed in this work improving statistical power. Indeed, PVM average rates were numerically higher than IFM and PVM had greater rates in five of seven muscle fibers in our previous *in vivo* work (Willingham *et al*., 2019). Importantly, the lower driving force across the electron transport chain (ΔG_NADH_ – ΔG_membrane potential_) in PVM suggests an increased pathway conductance compared to IFM and SSM even if there are no differences in the rate of NADH flux across groups.

It is also important to note that these data are collected from skeletal muscle of mice, and differences between human and mouse skeletal muscle are muscle type and measurement specific (Jacobs *et al*., 2013). Despite this, mitochondria in both human and skeletal muscle have fiber-type specific networks (Callahan *et al*., 2014; Dahl *et al*., 2014; Bleck *et al*., 2018; Caffrey *et al*., 2019), similar shapes (Katti *et al*., 2022a), organelle interactions (Picard *et al*., 2012; Picard *et al*., 2013a; de Almeida *et al*., 2023), and subcellular distributions (Vincent *et al*., 2019). Therefore, we would expect the findings observed herein to extend to human skeletal muscle. We also did not assess mitochondria relative to their proximity to nuclei. While nuclei generally localize to the PVM regions in oxidative skeletal muscle fibers (Rothstein *et al*., 2005; Glancy *et al*., 2014), the lack of PVM regions and high nuclear content in glycolytic muscle fibers may result in specialized perinuclear mitochondrial pools in these cells. However, perinuclear regions were not present within glycolytic fibers assessed here, precluding analyses. Future studies investigating how perinuclear mitochondria compare to IFM, SSM, and/or PVM will be of great interest. Lastly, our functional measurements were limited to basal conditions here. However, we previously reported that NADH redox and flux in both PVM and the combination of IFM and SSM responded similarly to mitochondrial uncoupling, fasting, or muscle contraction *in vivo*. How additional metabolic or pathological challenges influence the dynamic energetic response of PVM, IFM, and SSM will be the target of future work.

## Conclusion

Here, we define three structurally and functionally distinct subcellular regions of mitochondria within oxidative skeletal muscle cells defined by their proximity to myofibrils, capillaries, and the sarcolemma. Paravascular mitochondria are located in the gaps between the myofibrils and the sarcolemma created by an embedded capillary and are larger, rounder, and more connected with other mitochondria. This provides the structural capacity to maintain a more activated electron transport chain and a higher mitochondrial membrane potential. Subsarcolemmal mitochondria are found underneath the sarcolemma but away from a capillary. These mitochondria have a similar volume and surface area to volume ratio as the PVM, but functionally are more similar to the IFM, with a less activated electron transport chain and the lowest mitochondrial membrane potential. Interfibrillar mitochondria are found in the middle of the skeletal muscle fiber, are longer, and have more interaction with the sarcoplasmic reticulum. These mitochondria have a lower electron transport chain activation compared to PVM, but a slightly higher mitochondria membrane potential compared to SSM. Overall, these data suggest that skeletal muscle mitochondrial proximity to capillaries drives larger changes in mitochondrial structure and function than proximity to the sarcolemma. As such, future subcellular classifications of skeletal muscle mitochondria should aim to specify mitochondrial proximity to capillaries in addition to the sarcolemma and myofibrils.

